# Pre-stimulus oscillatory activity predicts successful episodic encoding for both young and older adults

**DOI:** 10.1101/358721

**Authors:** Jon Strunk, Audrey Duarte

## Abstract

Healthy aging is associated with declines in episodic memory performance that are due in part to deficits in encoding. Emerging results from young adult studies suggest that the neural activity during the time-period preceding stimulus presentation is sensitive to episodic memory performance. It is unknown whether age-related declines in episodic memory are due solely to changes in the recruitment of processes elicited by stimuli during encoding or also in processes recruited in anticipation of these stimuli. Here, we recorded oscillatory EEG while young and old participants encoded visual and auditory words that were preceded by cues indicating the stimulus modality. Alpha oscillatory activity preceding and following stimulus onset was predictive of subsequent memory accuracy similarly across age. Frontal beta oscillations linked to semantic elaboration during encoding were reduced by age. Post-stimulus theta power was positively predictive of episodic memory accuracy for old but not young adults, potentially reflecting older adults’ tendency to self-generate associations during encoding. Collectively, these results suggest that the preparatory mobilization of neural processes prior to encoding that benefits episodic memory performance is not affected by age.

## Introduction

Healthy aging is commonly associated with declines in episodic memory performance (Craik & Rose, 2012; Spencer & Raz, 1995). These declines are due in part to deficits in organization and/or binding of episodic information during encoding (Glisky, Rubin, & Davidson, 2001; Johnson, 1996; Old & Naveh-Benjamin, 2008). Encoding activity is typically measured as the difference in neural signal between subsequently forgotten and subsequently remembered events (Paller, Kutas, & Mayes, 1987; for review: Paller & Wagner, 2002). Numerous event-related potential (ERP) studies have confirmed that age-related changes in encoding contribute to older adults’ memory impairments (for review: Friedman, 2000).

Emerging results from young adult studies suggest that the neural activity during the time-period preceding stimulus presentation is also sensitive to episodic memory performance. The majority of these studies have investigated ‘pre-stimulus subsequent memory effects’ (preSME) ERPs during encoding (Cohen et al., 2015; Galli, Choy, & Otten, 2012; Gruber & Otten, 2010; Otten, Quayle, Akram, Ditewig, & Rugg, 2006; Otten, Quayle, & Puvaneswaran, 2010; Padovani, Koenig, Brandeis, & Perrig, 2011). Although the mechanisms underlying the preSME are not entirely clear, existing evidence suggests that it is highly dependent on the same factors that influence encoding and retrieval effects like stimulus modality (Golby et al., 2001; Wagner et al., 1998), type of encoding strategy implemented (Otten & Rugg, 2001; Paller et al., 1987), reward incentives (Adcock, Thangavel, Whitfield-Gabrieli, Knutson, & Gabrieli, 2006), and the format in which memory is tested (Paller & Wagner, 2002; Wagner, 1998). Consequently, pre-stimulus memory effects are thought to reflect, at least in part, preparatory mobilization of material specific and domain general processes that contribute to memory performance (Adcock et al., 2006; Addante, de Chastelaine, & Rugg, 2015; Otten et al., 2006; Xia, Galli, & Otten, 2018). It is unknown whether age-related declines in episodic memory are due solely to changes in the recruitment of processes elicited by stimuli during encoding or also in processes recruited in anticipation of these stimuli.

The majority of episodic memory studies have investigated ERPs, but scalp EEG, intracranial EEG and MEG studies have shown that oscillatory activity in several frequency bands both prior to and following event onset contributes to episodic memory performance. Unlike ERPs, oscillatory EEG responses are sensitive to underlying functional network dynamics with synchronization and desynchronization reflecting coupling and uncoupling of networks, respectively (for review: Duzel, Penny, & Burgess, 2010; Klimesch, Sauseng, & Hanslmayr, 2007). Both human and rodent research suggests that synchronous oscillations in the theta frequency band (4 – 8 Hz) reflect interactions between the hippocampus and cortical areas including the prefrontal cortex, which facilitate long-term memory (Klimesch, 1999; Nyhus & Curran, 2010). Mid-frontal theta power is typically greater for events that are remembered than those that are forgotten during encoding (Hanslmayr, Spitzer, & Bauml, 2009; Hsieh & Ranganath, 2014; Staudigl & Hanslmayr, 2013) and retrieval (Addante, Watrous, Yonelinas, Ekstrom, & Ranganath, 2011; Gruber, Watrous, Ekstrom, Ranganath, & Otten, 2013) particularly for events for which contextual associations are recollected (Addante et al., 2011; Gruber, Tsivilis, Giabbiconi, & Muller, 2008). Similar theta effects have been found preceding stimulus onset (Fell et al., 2011; Gruber et al., 2013; Guderian, Schott, Richardson-Klavehn, & Duzel, 2009). Because theta increases for remembered events have been shown for various kinds of stimuli and task conditions, it is likely that theta rhythms reflect domain general operations that contribute to episodic memory (Guderian et al., 2009).

Oscillations in the alpha (8 – 12 Hz) and beta (14 – 30 Hz) bands have also been found to relate to memory performance. In contrast to the increases in theta synchrony that contribute to successful encoding and retrieval, alpha and beta desynchronization following stimulus onset have been associated with memory success (for review: Hanslmayr, Staudigl, & Fellner, 2012; Sederberg, Kahana, Howard, Donner, & Madsen, 2003). Studies manipulating the strategies with which verbal materials were encoded have shown that decreases in beta power compared to baseline over anterior scalp sites may be related to semantic encoding rather than shallow or other forms of elaborative processing (Fellner, Bauml, & Hanslmayr, 2013; Hanslmayr et al., 2009). Intracranial evidence suggests that the left inferior frontal gyrus is a major generator of this beta encoding effects (Sederberg et al., 2003) and TMS evidence suggests that this desyncyhronization causally contributes to successful encoding (Hanslmayr, Matuschek, & Fellner, 2014). Alpha/beta desynchronization is believed to reflect processing within specialized neocortical areas sensitive to presented information (i.e. words, images, sounds, etc.) (for review: Hanslmayr, Staresina, & Bowman, 2016; Klimesch, 1999; Klimesch, 2012; Klimesch, Doppelmayr, Schwaiger, Auinger, & Winkler, 1999). Consistent with this idea is evidence showing that it varies topographically according to the nature of the materials associated with prior memories (Khader & Rösler, 2011; Waldhauser, Braun, & Hanslmayr, 2016; Waldhauser, Johansson, & Hanslmayr, 2012). As recollection is believed to depend, in part, upon neural reactivation of sensory information experienced during prior encoding (Rugg, Johnson, Park, & Uncapher, 2008), it follows that alpha and beta desynchronization during encoding may reflect processes that ultimately support recollection.

Given the multiple processes that these oscillations reflect, and the fact that they can be measured both prior to and following stimulus onset during encoding and retrieval, research in this area may be hugely important for elucidating the cause/s of age-related memory impairments. Very little work has been done to study the effects of aging on episodic memory with neural oscillations. Some EEG evidence from a visuospatial associative encoding task suggests that age-related reduction in theta synchronization following event onset may contribute to older adults’ memory impairments (Crespo-Garcia, Cantero, & Atienza, 2012). Similarly, MEG evidence suggests that increased stimulus-related theta power preceding encoding predicts relational binding success for the young but not the old in a short-term memory task (Rondina et al., 2015). These results are consistent with findings from short-term memory tasks showing age-related decreases in theta synchronization (Kardos, Toth, Boha, File, & Molnar, 2014; Karrasch, Laine, Rapinoja, & Krause, 2004). It is important to note that these studies assessed stimulus-induced changes in oscillatory power relative to baseline but did not compare oscillatory power for successful and unsuccessful memory trials. Thus, it remains unclear how aging affects pre- and post-stimulus oscillations that reflect encoding success, per se in episodic memory.

In the current study, we investigated age related differences in neural oscillations associated with episodic memory success both preceding and following event onset. Using a similar paradigm to that from Otten et al. (2010), we instructed young and old participants to make a semantic decision about audio and visual word stimuli during encoding. Modality congruent visual and auditory cues were presented prior to word onset at study and test to indicate the upcoming stimulus modality of the to-be-encoded or retrieved item. The majority of episodic memory studies investigating neural oscillations have been limited to visual events. Thus, the use of modality cues and stimuli allows us to determine whether pre-and post-stimulus oscillations reflect the engagement of domain-general processes or domain-specific perceptual processes that contribute to memory performance. Participants’ recognition memory was tested by intermixing previously presented items with new items and asking for their confidence that the item was previously presented. We predicted that greater theta synchronization and alpha/beta desynchronization prior to and following stimulus onset will be related to successful encoding. As alpha/beta desynchronization is believed to reflect processing within specialized neocortical modules (i.e. visual, auditory, semantic, etc.), we predict that domain-specific subsequent memory would be most likely within this frequency range. An age-related reduction in mid-frontal theta synchronization related to episodic memory success across stimulus categories would be consistent with previous reports of age-related reductions in hippocampal contributions to encoding and retrieval (Dennis & Cabeza, 2008; Dennis, Daselaar, & Cabeza, 2007; Dennis et al., 2008) and in hippocampal-PFC resting functional connectivity associated with worse memory performance (Andrews-Hanna, Saxe, & Yarkoni, 2014). An open question is whether aging impacts the preparatory mobilization of domain-specific perceptual processes and domain-general encoding mechanisms. Indeed, it is possible that at least some previous evidence of age-related differences in subsequent memory effects may be more accurately characterized as occurring or on-setting prior to stimulus presentation.

## Methods

### Participants

Participants were recruited from the Georgia Institute of Technology and the surrounding community. Thirty-four young adults participated for pay or course credit. Thirty-one older adults participated for pay. All compensation was paid at a rate of $10 per hour for each hour of participation. All participants were right-handed. Participants with neurological conditions, including Alzheimer’s disease, stroke, ADHD, untreated depression, schizophrenia, and epilepsy were excluded. All participants signed an Institutional Review Board (IRB) approved consent form prior to participation. All older participants completed on the Montreal Cognitive Assessment (MoCA) at the beginning of the experimental session to screen out possible mild cognitive impairment (Nasreddine et al., 2005). Participants were excluded if: they scored below age-adjusted norms on neuropsychological tests (Old: 1), did not complete the experiment (Young: 1; Old: 4), if memory performance was 2.5 standard deviations below the group mean (Young: 3; Old: 2), or if they had less than 12 artifact-free EEG epochs for a condition of interest (Young: 6; Old: 1). After exclusion, 24 young (10 males; Age: *M* = 21.37, *SD* = 3.04, Range = 18 – 32; Years Edu: *M* = 14.63, *SD* = 1.53, Range: 12 – 18) and 23 older (10 males; Age: *M* = 67, *SD* = 4.5, Range: 60 – 78; Years Edu: *M* = 16.26, *SD* = 2.34, Range: 12 – 21) adults were included in the analysis. Older adults had significantly more years of education than younger adults [*t*(45) = 2.87, p = 0.003].

### Stimuli

A pool of 480 concrete object nouns was used to create the study and test lists. Approximately half of each list consisted of items conceptually bigger or smaller than a standard computer monitor. The nouns were selected from the MRC Psycholinguistic Database (Wilson, 1988) with a written frequency of 10 – 50 occurrences per million (Kučera & Francis, 1967), a length of 3 – 12 letters, concrete range of 350 – 700, and image ability range of 500 – 700 (Coltheart, 1981). If multiple nouns had the same phonetic representation (e.g. “mail”, “male”), only one was retained. Each noun had an equal likelihood of being in the study or test list, as well as an equal likelihood of being presented as an auditory or visual item. Auditory stimuli were created with the software program Audacity (http://audacity.sourceforge.net/). All words were recorded by the same female voice and normalized (Mean Duration = 592 milliseconds (ms); Range = 250 – 1120 ms). All visual presentation occurred on a black background. Visually presented items were displayed in the center of the screen for 590-ms with white letters (Helvetica font, size 36). A white fixation cross was present on the screen at all times except during the period of visual cue and word presentation. The visual cue consisted of the fixation cross turning red for 250-ms, and the auditory cue was a 500Hz tone presented for 250-ms.

### Procedure

The experiment consisted of three parts: (a) incidental study phase (b) 30 minute delay, and (c) surprise recognition test. During the delay, participants completed the AX variant of the Continuous Performance Task (Braver et al., 2001) (not reported here). All participants received a short practice before each respective part of the experiment. Practice trials continued for each participant until they fully understood the task. An example of the trial structure, for the study and test period, is presented in Figure 1. The study period consisted of four blocks with 60 trials each. Each block contained an equal number of stimuli from each modality. Trials were pseudo-randomized with the requirement that the stimulus modality switch after a maximum of four trials. There were equivalent numbers of stay and switch trials in the experiment. Each trial began with a fixation cross randomly jittered between 300ms and 700ms by intervals of 50ms. Jitter was included in order to reduce expectancy related activity, like the contingent negative variation (CNV) prior to the cue onset. Participants were instructed to use the cue to prepare for the upcoming trial but were given no specific preparation instructions. For each trial, the cue always indicated the upcoming presentation modality of the word. For each word, participants decided if the word’s real-world referent was bigger or smaller than a standard computer monitor. Participants held a small USB number pad with both hands and pressed one button for “yes” and another for “no” using their thumbs. A one-second delay followed the participant’s response before the start of the next trial. If no response was made within two and a half seconds, the trial continued to the next trial.

**Figure 1:**
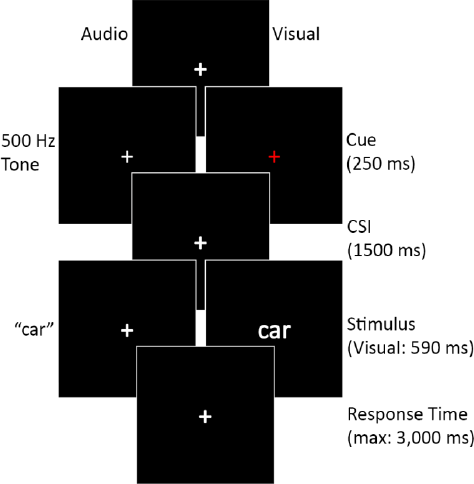
Trial structure and timing for both encoding and retrieval tasks.

The testing stage procedure used similar timing as the study phase, with the exception of the judgments made. The test period consisted of all 480 items (240 from the study list, and 240 new). Trials were pseudorandomized with the requirement that stimulus modality and old/ new status change after a maximum of four trials. There were equivalent numbers of stay and switch trials in the experiment. Each block contained an equal number of old visual, old auditory, new visual and new auditory, along with equal items from each bigger/smaller list. Each studied item presented during test was in the same modality as it was during study. For each item, the participant made an old/new decision with the following response options: “Old High Confidence”, “Old Low Confidence”, “New Low Confidence”, and “New High Confidence”. A fifth response option was available when participants were unsure of how to respond, to avoid guesses contaminating the other response categories. The trial proceeded one second after the subject response or, if no response, after four and a half seconds. Participants responded by pressing one of five keys on a number pad using the thumbs of both hands. Old and new judgments were counterbalanced between hands across participants.

### EEG Recording and Preprocessing

Continuous scalp-recorded EEG data was collected from 32 Ag-AgCl electrodes using an ActiveTwo amplifier system (BioSemi, Amsterdam, Netherlands). Electrode position follows the extended 10-20 system (Nuwer et al., 1999). Electrode positions included: AF3, AF4, FC1, FC2, FC5, FC6, FP1, FP2, F7, F3, Fz, F4, F8, C3, Cz, C4, CP1, CP2, CP5, CP6, P7, PO3, PO4, P3, Pz, P4, P8, T7, T8, O1, Oz, and O2. External left and right mastoid electrodes were used for referencing offline. Two electrodes placed superior and inferior to the right eye recorded vertical electrooculogram (VEOG), and two additional electrodes recorded horizontal electrooculogram (HEOG) at the lateral canthi of the left and right eyes. The ActiveTwo system replaces the traditional reference with a Common Mode Sense (CMS) active electrode and the ground with a Driven Right Leg (DRL) passive electrode. EEG was sampled at 1024 Hz with 24 bit resolution without high or low pass filtering.

Offline analysis of the EEG data was done in MATLAB 2015b with the EEGLAB (Delorme & Makeig, 2004), ERPLAB (Lopez-Calderon & Luck, 2014), and FIELDTRIP (Oostenveld, Fries, Maris, & Schoffelen, 2011) toolboxes. The continuous EEG data was down sampled to 256 Hz, bandpass filtered between 0.5 and 125 Hz, and then referenced to the average of the left and right mastoid electrodes. The continuous data was then epoched from 1500 ms pre-cue to 4000 ms post cue (i.e. 2250 ms post stimulus). In order to investigate slower frequencies, like theta, longer epochs are needed to account for loss of signal at each end of the epoch due to wavelet decomposition. The epoch range of interest was set to 600 ms pre-cue to 3400 ms post-cue (i.e. 1650 ms post-stimulus) in order to investigate oscillatory activity in both the pre- and post-stimulus intervals. In the time domain, each epoch was baseline corrected to the average activation across the whole epoch before artifact rejection and independent component analysis (ICA). ICA was run on the head electrodes in order to identify components reflecting blinks and eye movements, which were then removed from the data (Bell & Sejnowski, 1995; Delorme, Sejnowski, & Makeig, 2007; Hoffmann & Falkenstein, 2008). Each subject was manually inspected and any remaining epochs containing artifacts, such as muscle activity or saturation, were removed manually. Finally, each epoch was rebaseline corrected in the time domain to the average activity in the −600 to −100 ms time range (pre-cue onset).

### Time-Frequency Analyses

Each epoch was transformed into a time frequency representation using Morlet wavelets (Percival & Walden, 1993) with 5 cycles in 1Hz intervals between 2 and 30 Hz. After the time-frequency transformation, each epoch was reduced to the time range of interest (−600 to 3400 ms). Since each data point represents the weighted sum of the surrounding data points each epoch was down sampled to 50.25 Hz (Cohen, 2014). Then, individual subject averages were created for each condition and frequency of interest (Theta: 3 to 7 Hz, Alpha: 8 to 12 Hz, and Beta: 16 to 26 Hz) using a 10% trimmed mean (Wilcox & Keselman, 2003). Within each frequency, on the condition specific averages, a baseline normalization using relative change (Cohen, 2014) was calculated using the average frequency power over the −500 to −200 ms precue time range.

### Behavioral Analysis

Study or test items with a response faster than 200-ms, no response, or multiple responses were removed from further analysis. Memory accuracy was assessed using Pr (Snodgrass & Corwin, 1988). Pr (Hits – False Alarms) takes into account an individual subject’s false alarm rate (misclassifying a new item as an old item), which makes the ‘at chance’ rate equal to zero. Due to the subjective nature of the encoding task, accuracy was not assessed, although average subject agreement with predefined (Big/Small) word lists was .848 (*SD* = .114).

### EEG Analysis

In order to assess subsequent memory effects during encoding, we compared oscillatory activity between items subsequently recognized with high confidence (High Confidence Hits) and the combination of items subsequently recognized with low confidence and those misidentified as new. We refer to this combined category as Forgotten trials (for similar approaches: Hanslmayr et al., 2009; Otten & Rugg, 2001). It was necessary to combine low confidence hits with misses due to insufficient numbers of trials (<12) in these categories for many participants. Trial numbers across all participants for the conditions of interest are as follows: Visual High Confident Hits (mean = 62, SD=18), Audio High Confident Hits (mean=58, SD=17), Visual Forgotten (mean=41, SD=17), Audio Forgotten (mean=45, SD=17).

Significance testing was performed using Monte Carlo permutation tests with temporal and spatial clustering across our frequency bands of interest with the FIELDTRIP toolbox (Blair & Karniski, 1993; Maris & Oostenveld, 2007). Briefly, this method creates a reference distribution of t-statistics by using the t-statistics from randomizing the epochs across our conditions of interest with 2000 iterations to identify spatiotemporal clusters where the data in our conditions have reliably different distributions. Only spatiotemporal clusters that were reliable for over 200 ms and with a minimum of two neighboring electrodes were identified for follow-up analyses and quantification (for similar approaches: Addante et al., 2011; Gruber et al., 2013; Hanslmayr et al., 2009; Pastotter, Schicker, Niedernhuber, & Bauml, 2011; Staudigl, Hanslmayr, & Bauml, 2010).

This procedure was applied to contrasts of interest across all participants as well each group separately. We contrasted High Confidence Hits with Forgotten trials for both visual and audio items. This procedure was used for both mean differences and correlations with memory accuracy. Once significant spatiotemporal clusters were identified, the average cluster activity, across electrodes in the cluster, was used for subsequent analyses. Correlations were compared between younger and older adults by transforming each group’s correlation coefficient into a Z-score with Fisher’s transformation and followed by a two-tailed Fisher’s exact test with a 0.05 alpha level.

## Results

### Behavioral results

Response proportions for studied and unstudied items as a function of memory performance are listed in Table 1. Corrected recognition (Pr), collapsed across confidence, was calculated for both young (Visual: *M* = 0.541, *SD* = 0.180; Audio: *M* = 0.523, *SD* = 0.156) and older (Visual: *M* = 0.455, *SD* = 0.139; Audio: *M* = 0.397, *SD* = 0.134) adults. Accuracy was assessed with a 2 Modality (Visual, Auditory) X 2 Group (Young, Old) repeated measures ANOVA. The ANOVA revealed a main effect of Group [*F*(1,45) = 6.219, p = .016, η_p_^2^=0.121] and Modality [*F*(1,45) = 7.535, p = .008, η_p_^2^=0.147], but no interaction [*F*(1,45) = 1.976, p = .167, η_p_^2^=0.042]. These results show that older adults showed worse item recognition than the young and that memory was worse for audio items than visual items across groups. To parallel the EEG analysis, the same analysis was performed for High Confidence Pr estimates in the young (Visual: *M* = 0.533, *SD* = 0.165; Audio: *M* = 0.504, *SD* = 0.152) and older (Visual: *M* = 0.422, *SD* = 0.151; Audio: *M* = 0.371, *SD* = 0.141) adults. The results of the High Confidence Pranova revealed the same pattern as above. A main effect of Group [*F*(1,45) = 8.334, p = .006, η_p_^2^=0.156] and Modality [*F*(1,45) = 8.132, p = .007, η_p_^2^=0.153], but no interaction [*F*(1,45) = .670, p = .417, η_p_^2^=0.0.15]. Similar analyses of Low Confidence trials revealed near chance performance for both age groups. Consequently, we used High Confidence trials in all subsequent analyses.

**Table 1:** Recognition Memory: Proportion of Responses.

| Response: | *Old* |  |  |  | *New* |  |  |  |
| --- | --- | --- | --- | --- | --- | --- | --- | --- |
| Confidence: | *High* |  | *Low* |  | *High* |  | *Low* |  |
| Young Adults (n=24) |  |  |  |  |  |  |  |  |
| Old: Visual Items | 0.630 | [0.160] | 0.167 | [0.106] | 0.080 | [0.082] | 0.123 | [0.078] |
| Old: Audio Items | 0.609 | [0.136] | 0.169 | [0.098] | 0.085 | [0.085] | 0.137 | [0.073] |
| New: Visual Items | 0.097 | [0.066] | 0.158 | [0.116] | 0.382 | [0.233] | 0.363 | [0.177] |
| New: Audio Items | 0.105 | [0.066] | 0.151 | [0.095] | 0.371 | [0.202] | 0.374 | [0.173] |
| Older Adults (n=23) |  |  |  |  |  |  |  |  |
| Old: Visual Items | 0.567 | [0.159] | 0.146 | [0.107] | 0.163 | [0.121] | 0.123 | [0.072] |
| Old: Audio Items | 0.521 | [0.158] | 0.166 | [0.125] | 0.163 | [0.124] | 0.150 | [0.086] |
| New: Visual Items | 0.145 | [0.157] | 0.114 | [0.098] | 0.478 | [0.220] | 0.264 | [0.169] |
| New: Audio Items | 0.150 | [0.139] | 0.140 | [0.110] | 0.417 | [0.223] | 0.293 | [0.164] |

Note: Average Proportion [S.D.]

### Time-Frequency Results

At encoding we investigated the contrast between High Confidence Hit and Forgotten trials for mean differences (subsequent memory effects) as well as correlations with performance (high confidence Pr). We assessed our frequency bands of interest (theta, alpha, and beta) separately. Given the performance differences between visual and audio items, we chose to analyze the modalities separately. Using the cluster procedure described in the methods, we identified spatiotemporal clusters that showed significant mean differences or correlations with performance. Those clusters reaching significance are described below and quantified by a t-test on the average cluster power for mean differences. For correlations, the reported correlations correspond to the average cluster power correlated with the performance measure.

### Subsequent Memory Effects

#### Modality-general

No pre-stimulus or modality-specific average SMEs were observed in any frequency band. Cluster analysis across all participants did not reveal significant pre- or post-stimulus effects within the theta frequency band.

Within the alpha band, there was a significantly greater post-stimulus decrease in power for High Confidence Hits than Forgotten items across a cluster of 27 electrodes post-stimulus between 2,500 to 3,400 ms [t(46) = −3.859, p = 0.002]. This effect did not differ by modality [t(46) = −0.445, p=0.682], or group [t(45)= 0.943, p=0.338], as seen in Figure 2.

**Figure 2:**
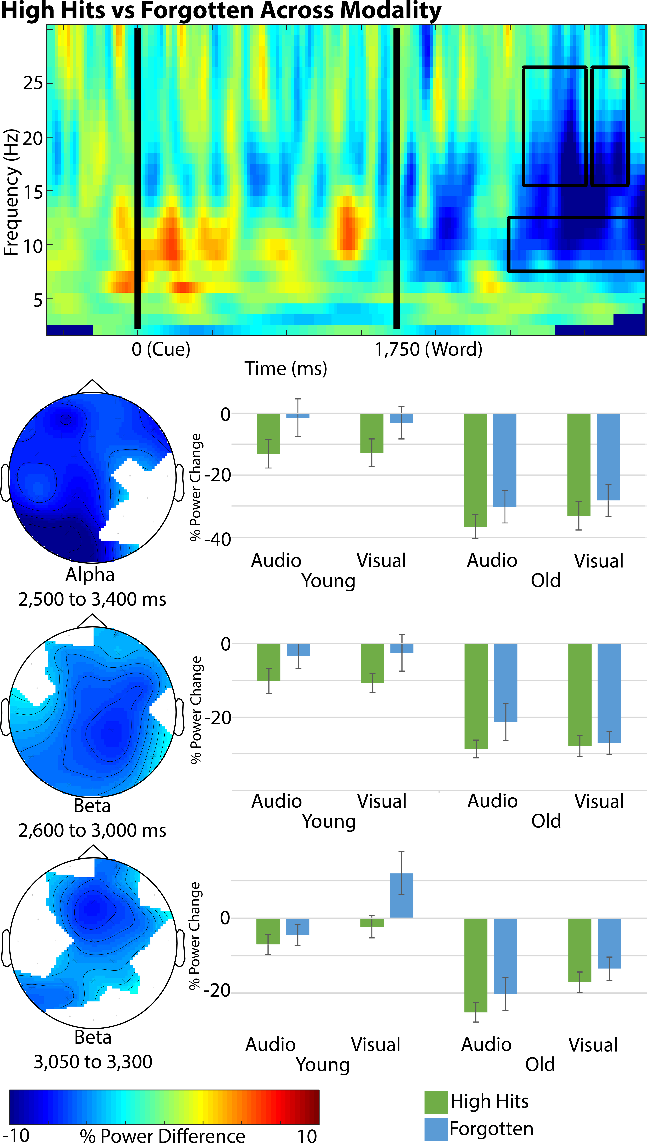
Heat map and topographic maps are averaged across all participants. Heat map represents the average of all significant overlapping electrodes across the alpha and beta frequency bands. Topographic maps only include the significant cluster electrodes and represent the difference in percent change from baseline between high confident hits and forgotten trials. Bar charts are the average percent change from baseline power within the electrode cluster. Error bars = 1 SEM.

Within the beta band, the analysis revealed a greater post-stimulus decrease for High Confidence Hits than Forgotten items across 28 electrodes post-stimulus between 2,600 to 3,000 ms [t(46) = −3.393, p = 0.001]. This effect did not differ by modality [t(46) = −0.831, p= 0.446], or group [t(45)= 0.834, p=0.429]. A second, later onsetting, cluster was identified within the beta band showing a greater post-stimulus decrease for High Confidence Hits than Forgotten items across 18 frontocentral electrodes post-stimulus between 3,050 to 3,300 ms [t(46) = 3.250, p = 0.001]. Follow-up analyses showed that the subsequent memory effect did not differ between modalities [t(46) = 1.624, p=0.113], or group [t(45) = −0.875, p = 0.385] but there was a marginally significant Modality x Group interaction [t(45) = 2.063, p = 0.056]. Follow-up tests showed that this was driven by a larger SME for visual [t(45) = 2.20, p = 0.035] but not audio trials [t(45) = 0.53, p = 0.627] for young than older adults, as seen in Figure 2.

### Subsequent memory effect correlations with memory performance

#### Modality-specific

For the alpha band, cluster analysis showed a significant negative correlation across all participants between High Confidence Audio Pr and the subsequent memory effect for audio trials across 11 bilateral centroposterior electrodes in the pre-stimulus time range between 150 to 1450 ms post-cue onset [r(45)=−0.41, p=0.004]. Fisher’s exact test showed no significant differences between the correlation magnitude between age groups [p = 0.226]. As can be seen in Figure 3, greater alpha desynchronization for hits than forgotten items prior to stimulus onset was predictive of better memory performance across age groups. Another significant negative correlation in the alpha band was found across 17 posterior electrodes between 1600 to 2800 ms post-cue (−150 to 1050 ms post-stimulus) [r(45)=−0.43, p=0.003], and Fisher’s exact did not find a significant difference between groups’ correlations [p =0.686]. These same clusters did not show reliable correlations between high confidence visual Pr and visual SME [r’s < 0.1, p’s > 0.528].

**Figure 3:**
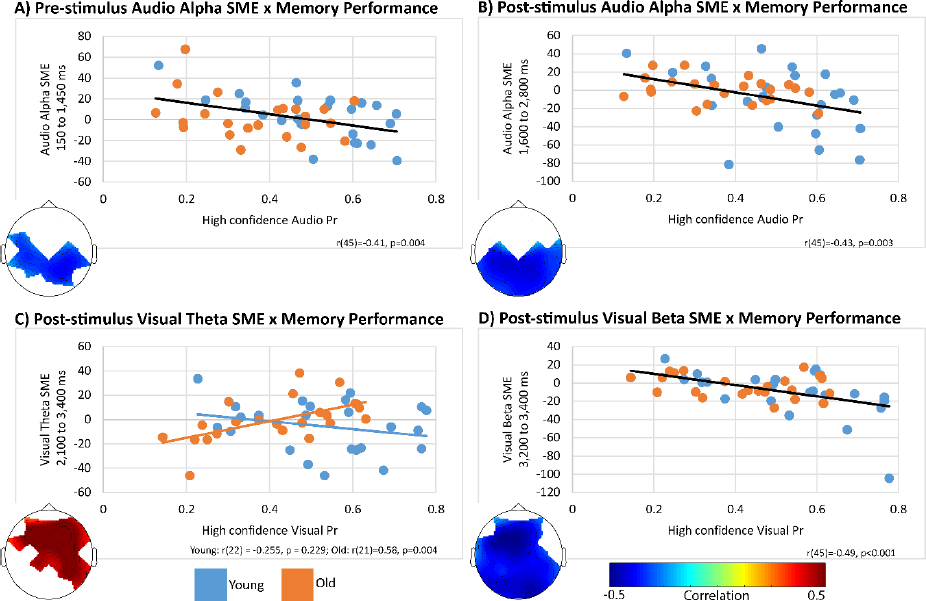
All plots are percent difference in power between high confident hits and forgotten trials. A) Prestimulus alpha power. B) Post stimulus alpha power C) Post stimulus theta power correlated with visual Pr in the old but not the young. D) Post-stimulus beta power.

For the beta band, a significant negative correlation was found between high confidence visual Pr and the subsequent memory effect for visual trials post-stimulus from 1450 to 1650 ms post-stimulus across 26 electrodes [r(45)=−0.49, p<0.001]. This correlation did not significantly differ between the young and older adults [p=0.278]. As can be seen in Figure 3, greater beta desynchronization for hits than forgotten items following stimulus onset late in the epoch was predictive of better memory performance for visual trials. This same cluster did not show reliable correlations between high confidence audio Pr and audio SME [r (45) = −0.03, p = 0.81].

Within the theta band, a significant positive correlation was found between high confidence visual Pr and the subsequent memory effect for visual trials post-stimulus between 2,100 to 3,400 ms across 16 fronto-central electrodes in older adults only [r(21)=0.58, p=0.004], young adults [r(22) = −0.255, p = 0.229]. Fisher’s exact test confirmed that the correlation coefficients between young and older adults were significantly different [p=0.003]. As can be seen in Figure 3, greater theta synchronization for hits than forgotten items following stimulus onset was predictive of better memory performance in older adults. This same cluster did not show reliable correlations between high confidence audio Pr and audio SME in either age group [r’s < 0.13, p’s > 0.53].

### EEG summary

Both modality specific and modality general effects were observed. Pre-stimulus subsequent memory effects over centroposterior electrodes were observed for both age groups but only for audio trials. Specifically, greater alpha desynchronization was correlated with better memory performance. This effect continued through the stimulus period. Several post-stimulus effects were observed: 1) a late on-setting and widely distributed beta desynchronization was predictive of better memory performance for visual trials only across age groups; 2) mid-frontal theta synchronization was predictive of better memory performance for visual trials in older adults only; 3) across groups and modalities, sustained and widely distributed alpha/beta desynchronization was greater for subsequently remembered items compared to forgotten items; 4) a late on-setting anterior beta desynchronization was greater for subsequently remembered than forgotten items across modalities. This effect was somewhat reduced in older adults.

## Discussion

Given accumulating research showing that the time-period before a to-be-remembered event influences the successful encoding of an event, we investigated neural oscillations both before (i.e. cue-stimulus interval) and after (post-stimulus interval) the to-be-remembered event during encoding. We investigated the extent to which age-related episodic memory impairments could be attributed to domain specific and domain general processes that support encoding both prior to and following stimulus onset. Behaviorally, we found young adults remembered more events than the older adults, and that memory was worse for audio items compared to visual items in both age groups. Post-stimulus, but not pre-stimulus memory effects, were affected by age with group differences for both material specific and material general oscillatory activity. Specifically, pre-stimulus centroposterior alpha desynchronization was greater for subsequent confident hits than forgotten auditorily presented words, continued through the stimulus period, and predictive of better memory accuracy across age. Post-stimulus beta desynchronization was greater for subsequent confident hits than forgotten visually presented words and predictive of better memory accuracy across age. Larger post-stimulus mid-frontal theta synchronization for visual trials was positively correlated with better memory performance in older adults only while frontal beta desynchronization was somewhat reduced for old relative to young adults. These results and their implications are discussed below.

### Behavioral results

Across age groups, memory accuracy for auditorily presented words was lower than that for visually presented words. This was not expected as a previous study using a design similar to our own showed no difference in memory performance between these stimulus modalities (Otten et al., 2010). Anecdotally, some participants, particularly older adults, commented that some auditory stimuli were more difficult to perceive than visual stimuli. One possible explanation for this difficulty is that the auditory stimuli were recorded in a female voice and aging is associated with loss of hearing for higher frequencies (Ferrand, 2002). If the auditory stimuli had been recorded in a male voice, with a deeper voice, we may have seen better memory performance for auditory items. Indeed, in the study by Otten and colleagues (2010), words were recorded in a male voice (Otten et al., 2010). Interestingly, we suspect that the perceived difficulty difference between visual and auditory trials may have contributed to at least some of the modality differences in the EEG oscillatory patterns, discussed below. Future research modulating stimulus quality within a single presentation modality would help directly answer this question.

### Pre-stimulus memory effects are sensitive to modality but not age

As discussed in the introduction, we wanted to determine whether anticipatory engagement of domain-specific perceptual and/or domain general encoding processes contribute to subsequent memory accuracy and the extent to which aging impacts this engagement. For auditorily presented words only, greater pre-stimulus alpha desynchronization for subsequent hits than forgotten items over centroposterior scalp sites was predictive of better memory accuracy for auditory events similarly across age. Alpha/beta desynchronization is believed to reflect the active engagement of specialized neocortical areas sensitive to processing the perceived materials (for review: Hanslmayr et al., 2016) Furthermore, anticipation of auditory stimuli has been associated with alpha oscillations over similar centroposterior scalp sites (Mazaheri et al., 2014). We suggest that the pre-stimulus memory effect observed here reflects the early activation of cortical areas that supports successful encoding of auditory events. The pattern of centroposterior alpha desynchronization persisted through the stimulus period supporting the idea that early engagement of the processes that support subsequent speech perception supports later memory performance for these events. These results are consistent with attention studies in which pre-stimulus cues facilitate early engagement of the domain-specific cortical areas engaged by the stimuli (for review: Driver & Frith, 2000; Luck, Chelazzi, Hillyard, & Desimone, 1997). Importantly, the lack of group difference suggests that this ability is not diminished by aging. Individuals, regardless of age, who mobilize these sensory processes early show superior memory performance.

It is not immediately clear why pre-stimulus memory effects were not also observed for visually presented words. Pre-stimulus domain-general effects were also not observed. Based on previous ERP evidence, we had predicted that pre-stimulus memory effects would be observed for both modalities (Gruber & Otten, 2010; Otten et al., 2006; Otten et al., 2010; Padovani et al., 2011; Padovani, Koenig, Eckstein, & Perrig, 2013). One possible explanation may relate to the different levels of difficulty between modalities. As discussed earlier, memory performance was worse for auditory than for visual items. It should be noted that alpha desynchronization is also modulated by memory load and effortful recovery of perceptual features (Lundqvist, Herman, & Lansner, 2011). Thus, the pre-stimulus activity pattern observed here together with the greater difficulty for the audio trials may suggest that anticipatory activation of speech processing areas was observed at least in part because these trials were more difficult to encode. One could argue that the centroposterior alpha desynchronization SME was not observed for visual trials not because it is auditory-related but because visual trials were easier to encode and placed lower demands on early mobilization of top-down attention. Although we cannot rule out this possibility, the lack of relationship between the magnitude of this effect and memory accuracy for visual trials, such that even low performing individuals showed no visual pre-SME, speaks against it. Collectively, these results support the idea that prestimulus memory effects are affected by numerous factors that may influence preparation including perceived difficulty (Speer, Jacoby, & Braver, 2003), reward (Gruber & Otten, 2010; Gruber et al., 2013), and the particular demands of the subsequent task (Otten et al., 2006; Padovani et al., 2011).

### Post-stimulus memory effects are sensitive to modality and age

Consistent with previous studies, widespread desynchronization in the alpha/beta frequency range was greater for subsequent hits than forgotten trials (Hanslmayr et al., 2009; Klimesch et al., 1996; Minarik, Berger, & Sauseng, 2018) between 900-1600 ms following stimulus onset. This SME was additionally insensitive to modality and age. The spatial distribution of this effect and the fact that it was similar across stimulus modalities suggest that it is likely supported by multiple neural generators and reflects several underlying processes that facilitate learning, including semantic elaboration and visual imagery. Specifically, given the verbal nature of the stimuli and the elaborative orienting task (“is this bigger than a monitor?”), it seems likely that semantic elaboration is one contributor to the SME effect. Similar subsequent memory effects in a similar time range within the beta band have been observed for verbal stimuli encoded in a deep but not shallow orienting task (Hanslmayr et al., 2009). TMS evidence suggests that this desyncyhronization causally contributes to successful encoding (Hanslmayr et al., 2014). Previous EEG evidence showing greater beta desynchronization for real than pseudo words, and intracranial data linking this activity to the left inferior frontal gyrus, suggest a role for beta oscillations in semantic processing (Hanslmayr et al., 2011). The similar spatial distribution and time course of the alpha SME shown here is consistent with data showing that alpha desynchronization may also support semantic processing (for review: Klimesch, 1999). Collectively, these findings together with the present results are consistent with the well-established idea that elaborative encoding facilitates episodic memory success (for review: Craik & Lockhart, 1972) and that older adults can effectively utilize this strategy to support episodic memory when instructed to do so (Naveh-Benjamin, Brav, & Levy, 2007). However, the beta desynchronization SME was somewhat reduced over frontal sites for old relative to young adults, particularly for visual trials late in the encoding epoch (1250-1550 ms). One possible explanation is that abbreviated semantic elaboration contributes to poorer encoding in older adults. Given the paucity of data investigating age-related changes in oscillatory EEG, future studies manipulating the orienting task demands will be necessary to draw more definitive conclusions regarding these group differences.

In addition to the modality invariant post-stimulus SMEs described above, there were also modality-specific correlations between post-stimulus activity and subsequent memory accuracy. A widespread beta desynchronization was greater for subsequent confident hits than forgotten visually presented words and predictive of better memory accuracy across age. This correlation was observed relatively late during encoding and overlapped both spatially and temporally with the average SME, described above, that was modality and age invariant. Thus, although this correlation was specific to the visual modality, the cognitive operations underlying this effect also support successful encoding for the auditory modality. We believe it is most likely that individuals who engaged in continued semantic elaboration and visual imagery in order to make size judgments were more likely to remember visually presented words. As discussed above, the anticipatory engagement of perceptual processes for auditorily presented words prior to stimulus onset and continuing through the stimulus period may have reduced the need for protracted encoding. Future studies manipulating stimulus modality together with other factors that influence engagement of pre-stimulus activity, including reward (Gruber et al., 2013), will be helpful in evaluating this possibility.

Contrary to what one might predict based on some prior evidence (Crespo-Garcia et al., 2012; Rondina et al., 2015), there was no evidence that subsequent memory activity in the theta band was reduced by age. In fact, older, but not young adults showed greater mid-frontal theta power for subsequent confident hits than forgotten visual trials that was predictive of greater memory accuracy. We had predicted that this effect would be observed both preceding and following stimulus onset, consistent with previous findings in young adults (Addante et al., 2011; Fell et al., 2011; Gruber et al., 2013; Gruber et al., 2008; Guderian et al., 2009; Hanslmayr et al., 2009; Staudigl & Hanslmayr, 2013). Furthermore, the correlation between the magnitude of the theta SME and memory accuracy that we observed for older adults has been shown for young adults in previous studies (Gruber et al., 2013). When considering these findings, it is important to discuss differences between the memory tasks and memory measures cross studies. In the current study, we compared activity between words subsequently recognized with high confidence with those recognized with low confidence or not recognized. Although events that are recollected are typically associated with high confidence judgments, it is also possible for events recognized on the basis of familiarity, for which no episodic details are recollected, to be based on high confidence (Yonelinas, 1994). As we did not direct participants to only respond with high confidence if they could recover specific episodic details or assess memory for objective details (i.e. source, context) we cannot be certain that our EEG contrast nor our estimate of memory accuracy is sensitive to recollection, exclusively. Previous studies showing mid-frontal theta band subsequent memory effects have often assessed memory directly for episodic details including object-location associations (Rondina et al., 2015), subjective recollection (Gruber et al., 2008), or source memory accuracy (Addante et al., 2011). These results have been taken as consistent with computational models suggesting that theta oscillations, generated by the hippocampus, facilitate episodic memory via functional interactions between the hippocampus and the cortex (Duzel et al., 2010; Hasselmo & Eichenbaum, 2005; Nyhus & Curran, 2010); and with the well know critical role of the hippocampus in memory for episodic details (for review: Eichenbaum, Yonelinas, & Ranganath, 2007). While it is unlikely that the theta SME measured at the scalp directly reflects hippocampal oscillations, due to its subcortical location and closed electric field, the long-range hippocampal-cortical theta interactions effects would be measurable (for review: Hsieh & Ranganath, 2014; Nyhus & Curran, 2010). We believe the most likely explanation for the lack of theta SME prior to or following stimulus onset for young adults may be a consequence of the manner in which we assessed memory success.

If the memory measure used in the current study was not particularly sensitive to the processes supported by mid-frontal theta synchrony, what might explain the beneficial impact of theta synchronization on subsequent memory accuracy for older adults? One possible explanation is that older adults either generated a greater number of episodic associations during encoding and/or used these associations to support their item recognition decisions to a greater extent than did young adults. Such an explanation is consistent with the previously observed discrepancy between age-related declines in objective and subjective tests of recollection (Ciaramelli & Ghetti, 2007; Duarte, Henson, & Graham, 2008; Duarte, Ranganath, Trujillo, & Knight, 2006; Mark & Rugg, 1998). Specifically, age-related declines in source or context memory are common despite relatively intact subjective reports of recollection, such as age equivalency in “remember” that are often rich in detail (Gallo, Korthauer, McDonough, Teshale, & Johnson, 2011). Subjective recollection can be based on less differentiated information or self-generated associations (i.e. thoughts, feelings) that are not typically assessed. By contrast, objective recollection requires participants to successfully bind experimental associations (e.g. spatial location, color, and modality) to studied materials and recover those specific associations during retrieval. The greater dependence of objective recollection than subjective recollection on executive functioning, which is disrupted by age, contributes to older adults’ disproportionate impairment for objective memory tests (Duarte et al., 2008; Duarte et al., 2006). If older adults in the current study based their item recognition decisions on recollection of thoughts and feelings associated with the stimuli, which they are more likely to self-generate than young adults (Carstensen & Turk-Charles, 1994; Comblain, D’Argembeau, Van der Linden, & Aldenhoff, 2004; Hashtroudi, Johnson, & Chrosniak, 1990; Kensinger, Allard, & Krendl, 2014; Leshikar, Dulas, & Duarte, 2015), then the theta synchrony effect may be unsurprising. We predict that if we had assessed recollection objectively, young adults, to a greater extent than older adults, would show the positive association between theta synchrony and memory accuracy that was shown here for older adults. An interesting future study would be to compare the relationship between the oscillatory activity patterns observed here and subjective and objective recollection memory tests. Importantly, the present pattern of results supports the idea that hippocampal integrity and/or hippocampal-cortical communication is not necessarily negatively impacted by age (Duarte et al., 2008; Dulas & Duarte, 2012; Morcom, Li, & Rugg, 2007).

## Conclusion

To our knowledge, this is the first study to investigate the effect of age on brain activity preceding episodic memory encoding. Our results are consistent with the idea that brain activity preceding to-be-encoded events, similar to that engaged during encoding, is beneficial for subsequent memory performance and that this anticipatory activity is spared by age. Oscillatory patterns of activity were largely similar across age, as consistent with data showing that older adults can successfully encode when given effective encoding tasks. An age-related reduction in frontal beta oscillations linked to semantic elaboration during encoding may contribute to their reduced memory performance. A greater reliance on theta oscillations for old than young adults may reflect older adults’ greater tendency to self-generate associations that may support their episodic memory accuracy when tasks are not constrained to objective experimental associations. An interesting question for future research is whether inducing oscillations via brain stimulation in one or more frequency bands prior to or during encoding might reduce age-related reductions in episodic memory performance.

